# A Novel Multiplex qPCR Assay for Detection of *Plasmodium falciparum* with *Histidine-rich Protein 2 and 3 (pfhrp2 and pfhrp3)* Deletions in Polyclonal Infections

**DOI:** 10.1101/2020.01.31.928986

**Authors:** Lynn Grignard, Debbie Nolder, Nuno Sepúlveda, Araia Berhane, Selam Mihreteab, Robert Kaaya, Jody Phelan, Kara Moser, Donelly A. van Schalkwyk, Susana Campino, Jonathan B. Parr, Jonathan J. Juliano, Peter Chiodini, Jane Cunningham, Colin J. Sutherland, Chris Drakeley, Khalid B. Beshir

## Abstract

**Background:** Rapid diagnostic tests (RDTs) that detect the malaria antigen histidine-rich protein 2 (HRP2) are widely used in endemic areas globally to confirm *Plasmodium falciparum* infection in febrile patients. The emergence of parasites lacking the gene encoding HRP2 and escaping RDT detection threatens progress in malaria control and elimination. Many health facilities in malaria endemic countries are dependent on RDTs for diagnosis and some National Health Service hospitals without expert microscopists rely on them for diagnosis out of hours. It is vital to study the emergence and the extent of such parasites globally to guide diagnostic policy. Currently, verification of the presence of such parasites in a blood sample requires a series of PCR assays to confirm the presence of *P. falciparum* and in the absence of amplicons from *pfhrp2* and/or *pfhrp3*, which encodes a cross-reactive protein isoform. These tests have different limits of detection and many laboratories have reported difficulty in confirming the absence of *pfhrp2* and *pfhrp3* with certainty.

**Methods:** We developed and validated a novel and rapid multiplex real time quantitative (qPCR) assay to detect *pfhrp2, pfhrp3,* confirmatory parasite and human reference genes simultaneously. We also applied the assay to detect *pfhrp2* and *pfhrp3* deletion in 462 field samples from different endemic countries and UK travellers.

**Results:** The qPCR assay showed limit of detection and quantification of 0.76-1.5 parasites per μl. The amplification efficiency, coefficient of determination (R^2^) and slope for the genes were 96-1.07%, 0.96-0.98 and −3.375 2 to −3.416 respectively. The assay demonstrated diagnostic sensitivity of 100% (n=19, 95% CI= (82.3%; 100%)) and diagnostic specificity of 100% (n=31; 95% CI= (88.8%; 100%)) in detecting *pfhrp2* and *pfhrp3* in. In addition, the qPCR assay estimates *P. falciparum* parasite density and can detect *pfhrp2* and *pfhrp3* deletions masked in polyclonal infections. We report *pfhrp2* and *pfhrp3* deletions in parasite isolates from Kenya, Tanzania and in UK travellers.

**Conclusion:** The new qPCR assay is simple to use and offers significant advantages in speed and ease of interpretation. It is easily scalable to routine surveillance studies in countries where *P. falciparum* parasites lacking *pfhrp2* and *pfhrp3* are a threat to malaria control.

## Background

Malaria is caused by infecting protozoan parasites of the genus *Plasmodium*. *P. falciparum* continues to be the predominant species with an estimated global incidence of more than 2228 million cases and about 405,000 deaths reported in 2018 (1). Immunochromatographic rapid diagnostic tests (RDTs), which use membrane-bound antibodies to detect parasite proteins in finger-prick blood samples, play a crucial role in malaria control successes in disease endemic countries. Early diagnosis is critical to malaria elimination and eradication programs and RDT deployment is an important component of the strategy. As a result, the global availability and scale of use of RDTs has increased dramatically over the last 10 years (2). Most RDTs used worldwide detect *P. falciparum* histidine-rich protein 2 (pfHRP2) and/or *Plasmodium* lactate dehydrogenase (pLDH) antigens. Some studies have shown that at least some pfHRP2-based RDTs also detect *P. falciparum* histidine-rich protein 3 (pfHRP3) due to a shared antigenic epitope (2–5). In sub-Saharan Africa, which bears 90% of the global malaria burden, RDTs accounted for 74% of diagnostic testing among suspected malaria cases in 2015, and pfHRP2-based tests were the most widely used (2).

Parasites with *pfhrp2* and/or *pfhrp3* genes (*pfhrp2/3*) deletions were first observed in South America and increasing reports of false-negative RDT results due to these parasites have now emerged from selected region of Africa and Asia (6–8). In some countries, a high proportion of RDT false-negative results due to these gene deletions has led to the changes in national diagnostic guidelines (9, 10). However, before undertaking any drastic changes in diagnostic testing policies or deploying less sensitive, less heat stable RDTs that detect alternative antigens, malaria programs need, robust epidemiological data about local *pfhrp2/3* deletion prevalence. The World Health Organization has prioritized studies of these parasites and developed a protocol for *pfhrp2/3* deletion surveillance (11). However, confirmation of pfhrp2/3 deletions using current techniques is challenging and time consuming. Most studies of *pfhrp2* and *pfhrp3* deletions deploy conventional nested PCR (nPCR) amplification of several genes followed by gel-electrophoresis (12). In this genotyping approach, at least three independent genes are used to ascertain the quality of DNA and the presence of *P. falciparum* parasites to avoid unintentional misclassification of pfhrp2/3 deletions in samples with low-concentration or degraded DNA (13, 14). The nPCR approach requires several rounds of PCR for each gene and running the gel-electrophoresis for each PCR product. The nested-PCR genotyping approach is labour-intensive, time consuming and is prone to contamination, particularly when deployed in large-scale surveillance studies. The various nPCR methods used differ in limit of detection, and this can cause type I and type II errors. Further, performance of reported the *pfhrp2* and *pfhrp3* PCR is variable, with wide ranging limits of detection and the risk cross-reactivity in some assays. In addition, gel-electrophoresis approaches do not detect deletions masked in polyclonal infections (6).

In this study, we report the development of a multiplex qPCR assay which simultaneously detects DNA from the human host, a single-copy parasite house-keeping gene, and the *pfhrp2* and *pfhrp3* genes, including in polyclonal *P. falciparum* infections, in a single reaction. We report the validation and application of this novel method using DNA samples derived from DBS and whole blood of field isolates and clinical samples. We also deploy the assay to estimate *P. falciparum* parasite density to rule out low parasite density as a factor for false RDT negative results (5, 13) when microscopic data is unavailable or if it is not reliable.

## Methods

### *Plasmodium falciparum* laboratory strains

Initial validation of the qPCR assay was performed using culture-adapted laboratory isolates with different *pfhrp2* and *pfhrp3* status, 3D7 (wildtype, West Africa origin), Dd2 (*pfhrp2 deletion, Indochina origin*), HB3 (*pfhrp3* deletion, Honduras origin) were obtained from the Malaria Research Reference Reagent Repository (http://MR4.org). A culture-adapted isolate lacking both genes (3BD5, double deletion) was also obtained from Thomas Wellems (NIAID, US). Parasite cultures of 3D7, Dd2, HB3 and 3BD5 were tightly synchronized as ring stage trophozoites *in vivo* to simulate infected peripheral blood similar to previously used methods (15, 16). The WHO *P. falciparum* International standard (Pf INT), a reagent comprising lyophilised whole blood from a single hyperparasitaemic individual was obtained from NIBSC UK. Undiluted, this reagent represents a parasitaemia of 9.8% which is equivalent to 4.9 × 10^5^ parasites per μl (15, 17).

### Clinical and field DNA samples

Clinical validation and application of the qPCR assay was performed using a convenience of 462 DNA samples derived from both DBS and whole-blood samples. Study site, sample collection and other details including IRB approvals have already been published for the study in which Eritrean samples were collected (9, 10). Parasite DNA was isolated from fifty dried bloodspot (DBS) samples from suspected malaria patients in Eritrean; 299 samples from whole-blood collected in EDTA from symptomatic Tanzanian and Kenyan patients and anonymised 113 samples from UK malaria patients. Whole blood collected in EDTA from UK travellers with confirmed *P. falciparum* infections in 2018 was obtained from the Public Health England Malaria Reference Laboratory (MRL), London, UK. Samples from Kenya (Ahero) and Tanzania (Bagamoyo) were collected between Oct 2016 and Dec 2018 as part of a study of parasite clearance after treatment with artemisinin combination therapy. These samples from Kenya and Tanzania have aliquots of cryopreserved blood samples and were selected for *pfhrp2/3* deletion study to identify *pfhrp2/3*-deleted parasites for culture-adaptation.

### DNA Extraction

DNA was extracted from DBS from Eritrea and from whole blood from the MRL and from cultured laboratory isolates using a robotic DNA extraction system (Qiasymphony, QIAGEN, Germany), as previously described (17). For the clinical samples collected in Kenya and Tanzania, 200μl of whole-blood was extracted using the QIAamp Blood Mini Kit (Qiagen) into 200μl of Buffer EB as per the manufacturer’s instructions.

### Multiplex qPCR development

#### Gene target selection and primer design

To design highly specific amplification primers that are conserved across global *P. falciparum* isolates we carried out multiple alignment of *pfhrp2* gene sequences from 1581 published *P. falciparum* genomes (MalariaGEN) from Africa, SE Asia and South America, and a similar alignment was also carried out for *pfhrp3* (Figure S1). The DNA sequence of the genes were obtained from publicly available genomic data and the processing of the data has been described in our previous report (6). We have also aligned the 3D7 DNA sequence of *pfhrp2* (PF3D7_0831800) and *pfhrp3* (PF3D7_1372200) to ensure that the conserved primers of the two genes do not cross-bind and are specific to *pfhrp2* and *pfhrp3* respectively (Figure S2). The DNA sequences of *pfhrp2* and *pfhrp3* were aligned using Geneious v. 10 (Biomatters, USA) We used *Plasmodium falciparum* lactate dehydrogenase *(pfldh,* PF3D7_1324900), coded by a single-copy gene on chromosome 13, as a confirmatory gene for the presence and quality of parasite DNA as well as a target for measuring parasite density. We used previously published qPCR methods, with some modification of reaction conditions to amplify *pfldh* (13) and the human beta tubulin gene (*HumTuBB*) (Table S1)(18). The latter was used both as an internal control and as a normalizer for measurement of parasite density and for detection of *pfhrp2/3* deletion in polyclonal infections. All primers and probes were ordered from Eurofins Scientific (Germany).

#### Modification of pfhrp2 primer at the 3’ end

Due to limited availability of suitable conserved target sequences region in *pfhrp2* and *pfhrp3* that are dissimilar between the isoform genes and to prevent non-specific cross-binding of the *pfhrp2* primers to *pfhrp3*, we used a strategy altering nucleotides located at the 3’ end region (within the last 5 nucleotides) of both *pfhrp2* primers (Table S1). In total, we designed six primers with different modification at the 3’ end of the forward and reverse *pfhrp2* primers, which were then tested empirically to identify primer pairs that delivered the best specificity while maintaining product yield (sensitivity).

#### Assay optimization

The multiplex qPCR assay designed in this study was optimized for primer and probe hybridization temperature; different primer and probe concentrations and different MgCl_2_ concentrations. All the optimization analyses were performed in triplicates in a RGQ rotor-gene (Qiagen, Germany).

The optimized final reaction conditions were performed in a final volume of 25 μl containing 1.6X NH_4_ buffer (Bioline); 4 mM MgCl_2_ (Bioline); 800nM dNTPs (Bioline), 200nM of *pfhrp2* primers, 200nM *pfhrp3* primers, 120 nM of *pfldh*, 120 nM *HumTuBB* primers, 120 nM of *pfhrp2* probe, 120 nM *pfhrp3* probe, 80 nM of *pfldh* probe and 80 nM *HumTuBB* probe; 2 units of biotaq polymerase and 5 μl of extracted DNA. The optimal thermocycling conditions selected were 3 mins at 95°C, followed by 45 cycles of 15 sec at 95°C; 30 sec at 54°C and 30 sec at 72°C.

#### PCR efficiency, linear dynamic range and limit of detection

We conducted assay performance analysis based on the MIQE guidelines(19). The efficiency of primer and probe combinations for each gene and linear dynamics of the qPCR assays were evaluated using seven 4-fold dilutions of Pf INT (from 12500 to 3 parasites per μl). The last dilution series (3 parasites per μl) was then further diluted 2-fold (from 3 to 0.38 parasites per μl) to determine the measured limit of detection (LOD). The three laboratory strains (Dd2, HB3 and 3BD5) with known *pfhrp2/3* status were included in the evaluation linear dynamic of the linear range to examine the effect of DNA concentration on the cross reactivity of the primers.

#### Assay precision, analytical sensitivity and specificity

Performance of the multiplex qPCR *pfhrp2/3* assay was evaluated by measuring coefficient of variation across the seven four-fold dilution series of Pf INT. The specificity and sensitivity as well as the robustness of the assay was evaluated by testing the lowest two concentrations of Pf INT replicates of eight and 20 *P. falciparum*-negative whole blood samples in three different experiments.

#### Detection of parasites with pfhrp2 and pfhrp3 *deletion* hidden in polyclonal infections

To investigate whether the qPCR assay could detect *pfhrp2/3*-deleted parasites hidden in polyclonal infections robustly and accurately, we generated pairwise mixtures of known *P. falciparum* genotypes (Dd2 and Pf INT, HB3 and Pf INT, and 3BD5 and Pf INT) at different ratios i.e. 1:1, 5:1, 10:1, 100:1, 1000:1, 10,000:1, 1:100,000, 1:10,000, 1:1000, 1:100, 1:10, 1:5, 1:1. We estimated the abundance of *pfhrp2/3* deletion genotypes in the mix relative to the whole parasite biomass as measured by relative quantification normalized to *pfldh* (measures the DNA of all strains) and *humTuBB* genes. The qPCR assay was also used to detect *pfhrp2/3* deletions in patient samples with known polyclonal infection in patient samples, as determined by a peviously published high resolution melting qPCR assay (20).

#### Relative quantification using the pfldh gene

We used *pfldh* as a parasite target and *HumTuBB* as a normalizer to estimate relative parasite density of each sample. The lowest parasite density (parasite per μl) with a coefficient of variation of less than 35% was considered limit of quantification for the assay(21). We determined the performance of the *pfldh* qPCR in the multiplex assay by comparing parasite densities determined by a published *pgmet* duplex qPCR assay using samples from Eritrea (18).

#### Application to DNA from diverse field samples

The validated qPCR assay was then applied to DNA extracted from diverse field samples. Samples with Cq values lower than the Cq value of the limit of detection of individual gene are determined to be negative for the gene. Samples with *HumTuBB* positive and *pfldh* positive but negative for *pfhrp2* or *pfhrp3* are determined to be *pfhrp2*-deleted and *pfhrp3*-deleted respectively. Samples with *HumTuBB* positive but pfldh negative are determined to be parasite negative. Finally, samples with *HumTuBB* negative are considered to be invalid and DNA extraction and/or PCR experiment should be repeated.

### Data Analysis

For all amplification curve analyses, the quantification cycle (Cq) threshold was placed above the amplification curve of No Template Control (NTC) and any crossing point between the Cq threshold and the amplification curve was considered positive (Cq value) for the specific sample. To evaluate assay precision, we calculated the coefficient of variation (CV) of parasite density (parasite per μl) as follows: CV% = (standard deviation/mean) × 100. To determine the limit of detection (LOD), we calculated the percentage of positive samples and the lowest sample with more than 3 parasites per PCR and with ≥ 95% of replicate samples detected was considered LOD (19). Similarly, to determine the lower limit of quantification (LOQ), we calculated the parasite density using delta delta method and the sample with lowest parasite density (parasite per μl) with a CV ≤35% was considered LOQ for the assay (21). Throughout the manuscript CV refers to variation in parasite density (parasite per μl).

For estimation of relative abundance of *pfhrp2/3* deletion in a mixed-strain infection we used delta delta relative quantification method (18) as follows:

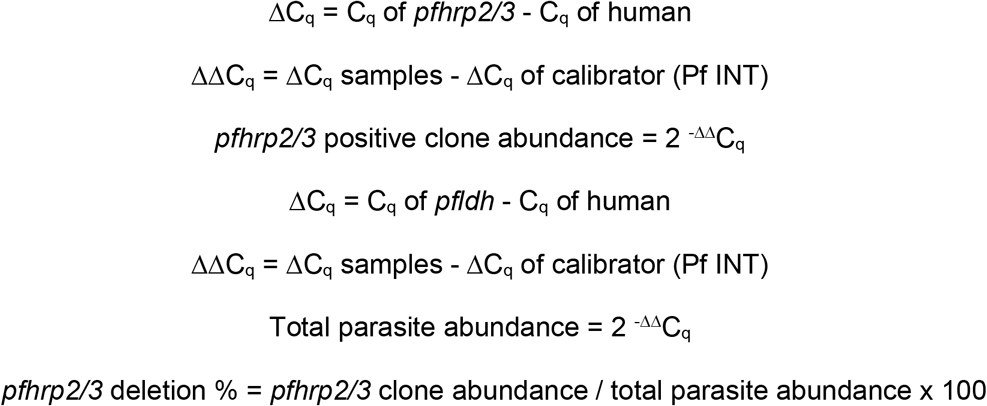

Before calculating the relative abundance of *pfhrp2/3* strains, the C_q_ threshold of the positive control (calibrator, Pf INT, 0.98% parasitaemia) was adjusted in each channel in such a way that the C_q_ value is similar in all parasite target genes.

For comparison of parasite density estimated by two different _q_PCR assays, we used STATA (v 15, USA) software to perform linear regression.

### Ethics approval and consent to participate

Ethical approval for collection of samples was obtained from each local ethical committee in Eritrea (Eritrean MOH Research and Ethical Committees), Kenya (KEMRI IRB, 3293) and Tanzania (MUHAS IRB, DA.282/298/01). Ethical approval for the samples from MRL patients was obtained from NHS England Research Ethics Committee (18/LO/0738). The ethical approval for the laboratory work for the Eritrean and MRL samples was obtained from LSHTM Ethical Review Committee (#11979 and #14710 respectively).

## Results

### Primer and probe selection, and *in silico* analysis

Initial assessment of *pfhrp3* primers across exons 1 and 2 showed cross reactivity with *pfhrp2* target (Figure S3) and a new set of primers within a specific conserved region of exon2 of *pfhrp3* was designed. After initial assessment *pfhrp2 and pfhrp3* primers for specificity and length of the probe, two sets of *pfhrp3* primers and three sets of *pfhrp2* primers, including primers with modifications at the 3’ end, were selected for testing (Table S1). For *pfhrp2* primers, of the nine combinations used, the lowest C_q_ value was obtained when *pfhrp2*_F1 and *pfhrp2*_R2 (modification at 3’ end) were combined and were selected for further optimization (Table S2). Since one single mutation in *pfhrp2* forward primer was found in one sample in The Gambia and three samples in Ghana we have nucleotide redundancy in the synthesis of *pfhrp2_*F1 primer to reflect these mutations (Table S2). The other *pfhrp2* primer combinations either produced fluorescence signal in Dd2 (*pfhrp2*-deleted laboratory strain) due to cross binding to *pfhrp3* or generated relatively higher (more unfavourable) C_q_ values in *pfhrp2*-positive lab strains (HB3 and Pf INT) compared to the selected primer combinations (Figure S4). Interestingly, a single nucleotide change (T to G) decreased the C_q_ value by 7 (30 to 23) while two nucleotide changes (T to G and T to G) decreased the C_q_ value by 2 (30 to 28) (Table S2).

### Detection of *pfhrp2* and *pfhrp3* in laboratory strains

We followed the MIQE guidelines for optimization of the qPCR assay; for analysis and reporting of the data (19). The analytical specificity and sensitivity of each newly designed primer and probe combination was validated in monoplex and multiplex qPCR assays on Dd2, HB3, 3BD5 and 3D7, laboratory strains and Pf INT. The assay correctly determined the *pfhrp2* and *pfhrp3* status of the four laboratory strains and the Pf INT. No amplification of either *pfhrp2* or *pfhrp3* was observed in Dd2 and HB3 respectively, while 3BD5 produced no fluorescence signal in either the *pfhrp2* or *pfhrp3* channels (Figure 1). 3D7 and Pf INT were *pfhrp2* and *pfhrp3* positive.

**Figure 1.**
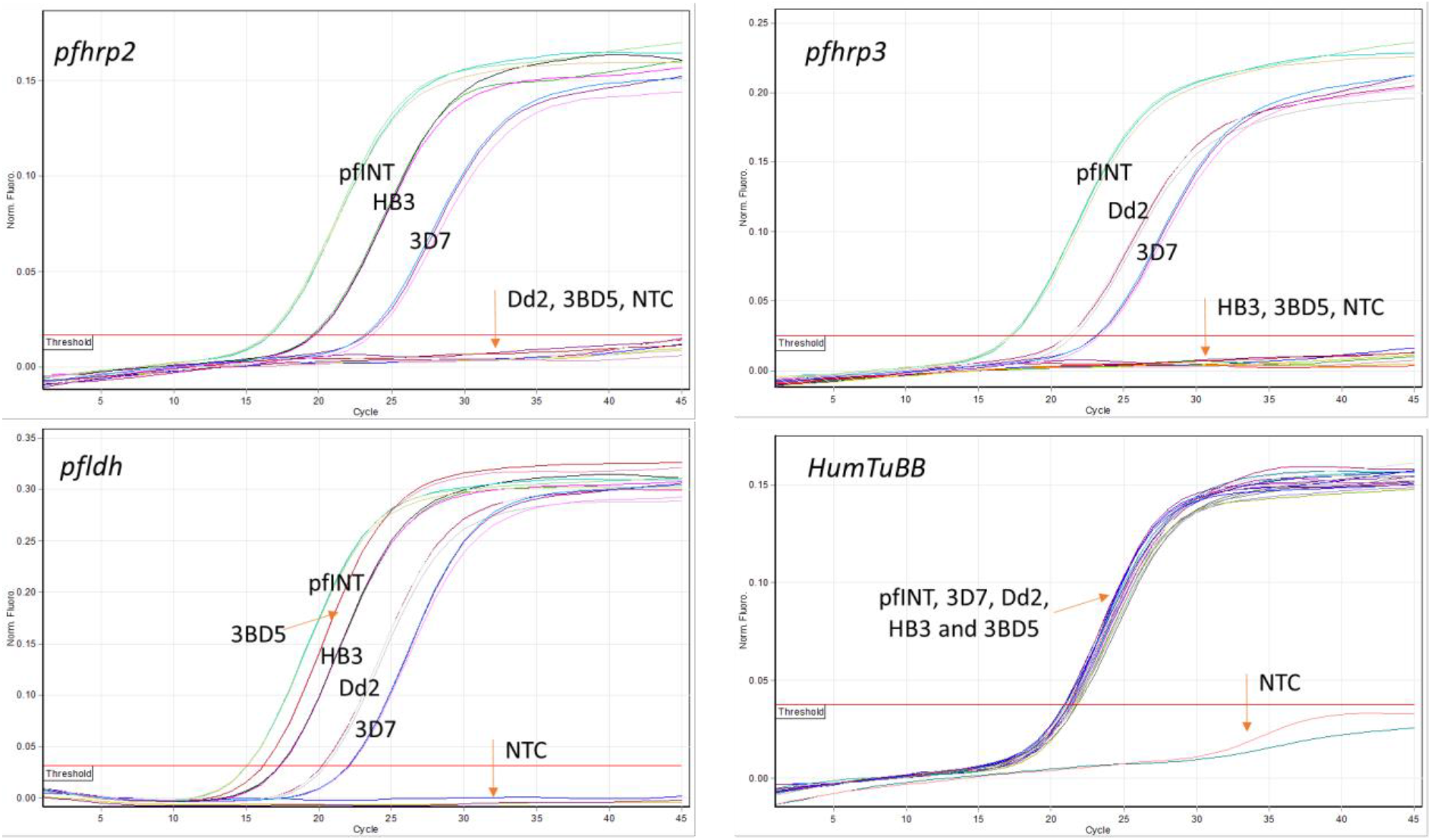
Amplification of four laboratory clones (3D7, Dd2, HB3 and 3BD5) and Pf INT in three parasite targets (*pfhrp2*,*pfhrp3* and *pfldh*) and a human beta tubulin gene (HumTuBB). Laboratory clones 3D7 (wild type), Dd2 (*pfhrp2* deletion), HB3 (*hrp3* deletion), 3BD5 (both *pfhrp2/3* deletion) and Pf INT (*Plasmodium falciparum* WHO International Standard, both *pfhrp2/3* present) were amplified in triplicate targeting four different genes; *pfhrp2*, *pfhrp3*, *pfldh* and *HumTuBB*. The red horizontal line marks the threshold, the normalised fluorescence is measured on the y-axis and the number of cycles on the x-axis. Orange arrows point to different clones and the negative no template control (NTC).

### Analytical sensitivity of the qPCR assay

The *pfhrp2*, *pfhrp3* and *pfldh* qPCR assays allowed detection of three parasites per μl with a C_q_ standard deviation (SD) of 0.58, 0.47, 0.41 respectively (Table 1A), which corresponds to parasite density CVs of 9.4%, 9.3% and 8.9% of respectively (Table 1B). The sample with dilution of the 1.5 parasite per μl showed a C_q_ SD of 0.80, 0.71 and 0.74, which corresponds to parasite density CVs of 32.6%, 26.1% and 28.1% respectively (Table S4). Though the assay also detected as low as 0.76 parasites per μl the SD value was very high (1.49, 1.37 and 1.48 respectively), and this corresponds to parasite density of CVs of 109%, 99% and 108% (Table S4). Therefore, the lowest parasite density that can be quantified with CV of 35% lies between 1.5 and 0.76 parasites per μl.

**Table 1A:** Precision of the parasite target genes: Mean, standard deviation (SD) of quantification cycle (C_q_) were calculated from amplifications of seven Pf INT 4-fold dilutions (in triplicate) with *pfhrp2*, *pfhrp3* and *pfldh* assays.

| Parasite density (p/$\mu\text{L}$) | Mean $C_q$ values | | | | | |
| --- | --- | --- | --- | --- | --- | --- |
|  | *pfhrp2* |  | *pfhrp3* |  | *pfl dh* |  |
|  | Mean | SD | Mean | SD | Mean | SD |
| **12500** | 23.18 | 0.06 | 23.46 | 0.02 | 22.44 | 0.04 |
| **3125** | 25.49 | 0.02 | 25.82 | 0.19 | 24.77 | 0.06 |
| **781** | 27.11 | 0.07 | 27.55 | 0.07 | 26.48 | 0.08 |
| **195** | 29.02 | 0.07 | 29.54 | 0.12 | 28.33 | 0.16 |
| **49** | 31.28 | 0.59 | 32.16 | 0.73 | 30.81 | 0.77 |
| **12** | 33.17 | 0.66 | 33.73 | 0.54 | 32.40 | 0.57 |
| **3** | 35.36 | 0.58 | 34.76 | 0.47 | 35.12 | 0.41 |
| **Total variance** | 28.95 | 0.15 | 29.31 | 0.18 | 28.32 | 0.17 |

**Table 1B:**
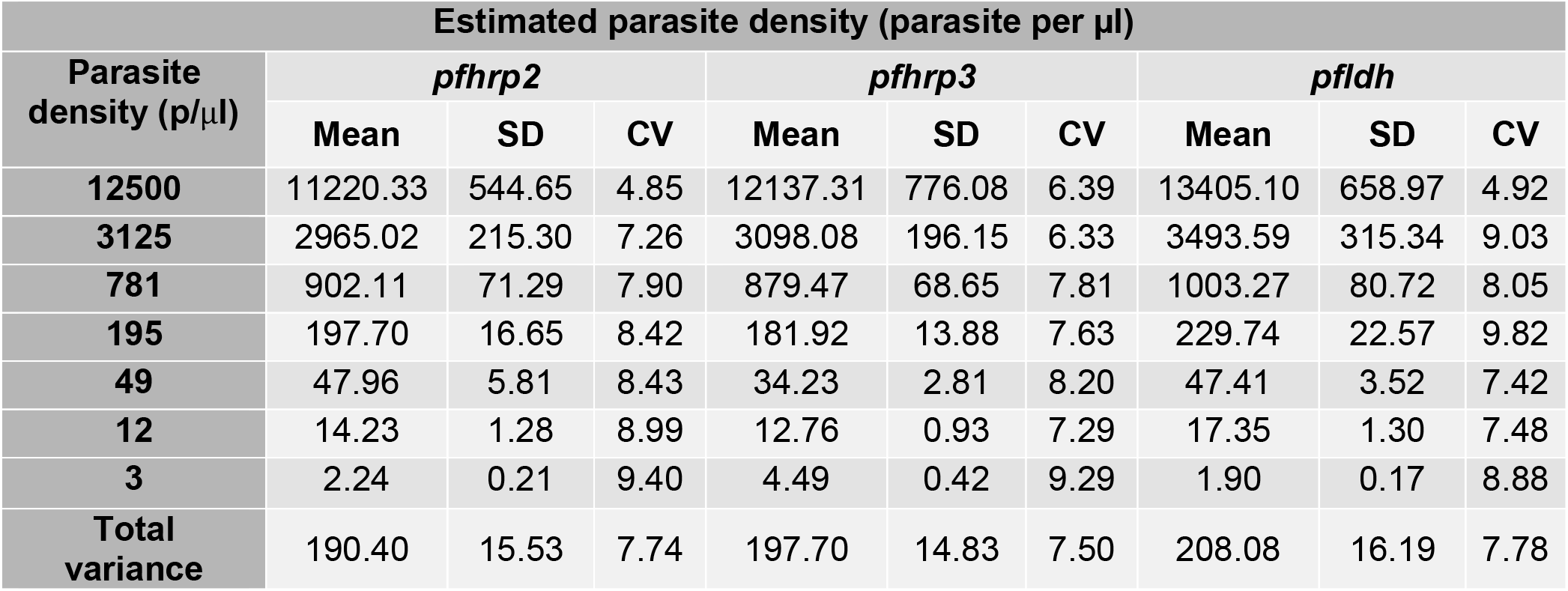
Precision of the parasite target genes: Mean, standard deviation (SD) of coefficient of variation (CV) parasite density were calculated from amplifications of seven Pf INT 4-fold dilutions (in triplicate) with *pfhrp2*, *pfhrp3* and *pfldh* assays.

### PCR efficiency and dynamic range

In order to measure abundance of *pfhrp2* and *pfhrp3* clones relative to *pfldh* and estimate parasite density relative to human DNA, the four targets should have similar amplification efficiency. The efficiency of each qPCR was evaluated on 4-fold serial dilutions of Pf INT (from 12500 to 3 parasite per μl) and each primer and probe combination resulted in similar PCR efficiency (Figure 2) and this was evident by the C_q_ value generated by each combination. A 4-fold dilution of the Pf INT produced linear standard plots with 98%, 96%, 98% and 1.07% PCR amplification efficiency for *pfhrp2*, *pfhrp3*, *pfldh* and *HumTub* primer pairs respectively (Figure 2). The coefficient of determination (R^2^) of the standard curve was 0.96-0.98 with a slope value of −3.375 to −3.416 for each assay.

**Figure 2.**
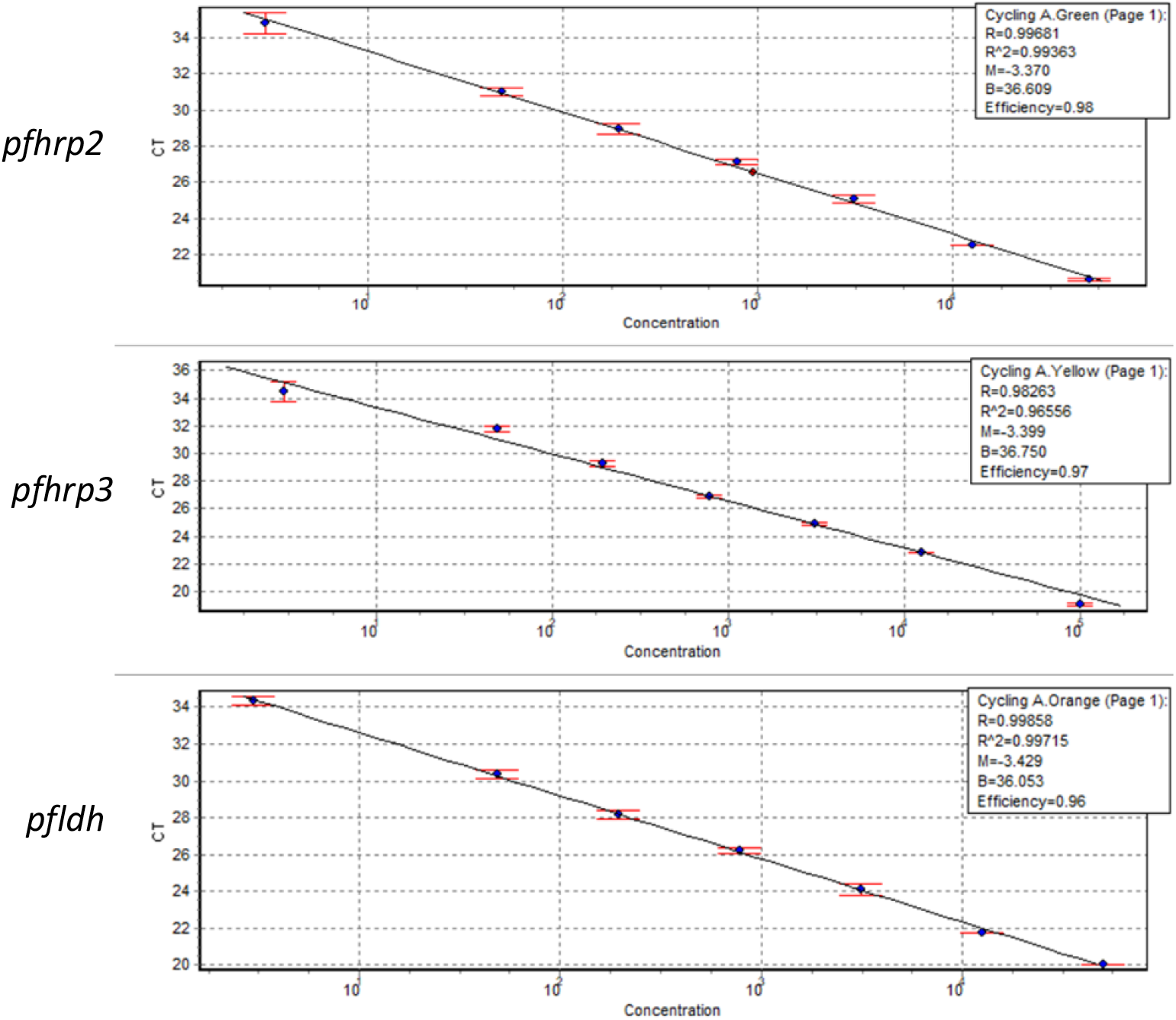
Standard curve of 7 Pf INT samples diluted 4-fold starting from 12500 parasites per μl. The three parasite assays (*pfhrp2*, *pfhrp3* and *pfldh*) detected as low as 3 parasites per microliter with amplification efficiency of 98%, 97% and 96% and coefficient of determination (R2) −3.37, −3.39 and −3.43 respectively.

### Performance of the qPCR assays

We assessed the precision of the assay by testing seven 4-fold dilution series of the Pf INT (starting concentration, 12500 parasites per μl) in triplicate on three separate occasions. The mean SD for cycle quantification across the seven samples was 0.15, 0.18 and 0.17 for *pfhrp2*, *pfhrp3* and *pfldh* target genes respectively (Table 1A), which corresponds to a calculated parasite density of CVs of 7.4%, 7.5% and 7.8% respectively (Table 1B). The sensitivity and specificity as well as the robustness of the assay was assessed using *P. falciparum*-negative blood samples and Pf INT (3 and 1.5 parasites per μl) in eight replicates on three different occasions. All the control samples were negative, while the Pf INT was positive indicating the absence of amplification inhibition and non-specific amplification (Table S4).

### Detecting parasites with *pfhrp2/3* deletions hidden in polyclonal infections

The ability of the qPCR assay to detect minor or major laboratory clones with *pfhrp2/3* deletion was assessed using artificially mixed laboratory strains Dd2, HB3, 3BD5 and Pf INT. Both *pfhrp2* and *pfhrp3* qPCR assays showed good performance in detecting minor (as low as 20%; CI, 7.94-27.05) and major (as high as 99.99%; CI, 99.99-100) *pfhrp2/3*-deleted clones in the artificially mixed laboratory clones (Table S4). The *pfhrp2/3* assays can also detect as low as 10% *pfhrp2/3* deleted clones but the value lies within the confidence interval of replicate experiment of a single clone sample and therefore lacks confidence.

### Workflow and throughput time

Performing the multiplex qPCR requires three steps: approximately 20 minutes for reaction setup takes ~20 minutes, two hours for runtime 30 minutes for analysis. Over all the estimated time was three hours for 72 reactions on the Rotorgene Q platform. In comparison, the estimated time for the conventional method, deploying several nested PCR assays in a 96-well plate format followed by electrophoresis is approximately 30 hours. Time for DNA extraction is the same for both.

### Validation and application of the qPCR assay on field samples

To determine the diagnostic sensitivity and specificity of the qPCR assay for detecting *pfhrp2/3* deletions in field samples and to provide population estimates of deletion prevalence, we first investigated 50 DNA samples obtained from confirmed *P. falciparum* patients from Eritrea whose *pfhrp2/3* deletions were previously determined using the conventional nPCR method (10). Results obtained from the qPCR assay were fully concordant with previously reported results from the same samples using the conventional nPCR method (10). The qPCR assay correctly detected all the *pfhrp2/3* positives (100% sensitivity for *pfhrp2*, n=19, 95% CI = (82.4%, 100%); 100% sensitivity for *pfhrp3*, 95% CI= (66.4%; 100%), n=9) and accurately determined the absence of *pfhrp2/3* in the remaining samples (100% specificity for *pfhrp2,* n=31, 95% CI=(88.8%,100%); 100% specificity for *pfhrp3*, n=41, 95% CI=(91.4%,100%)) (Table 2). We then assessed samples obtained from clinical malaria patients from the MRL (n=113), Kenya (n=150) and Tanzania (n=149). For the MRL samples, we selected 113 samples from countries with reported *pfhrp*2/3 deletion or high risk of emergence of *pfhrp*2/3 deletion (22). The countries include Eritrea (n=1), Ethiopia (n=2), Kenya (n=20), Tanzania (n=16), Uganda (n=27), Sudan (n=29), South Sudan (n=10), Djibouti (n=1), Somalia (3) and east Africa (unknown country, n=2). Analysis of the *pfhrp2/3* status of the MRL samples by qPCR showed 1.8%, 1.8% and 6.3% *pfhrp2−/3+*, *pfhrp2+/3−* and *pfhrp2−/3−* deletions respectively. One *pfhrp2* deletion occurred in a UK traveller from Sudan and five *pfhrp3* deletions occurred in UK travellers from Ethiopia (1), Sudan (1), South Sudan (2) and Uganda (1). There was evidence of *pfhrp2/3* deletions in polyclonal infections in 3.5% of MRL samples in total (Table 4). Of the 149 whole blood samples collected from Tanzania, 1 sample (0.7%) carried a *pfhrp2* deletions and another sample (0.7%) carried a *pfhrp3* deletion while no deletions were observed in the 150 Kenyan samples. However, there were 5 samples (3.4%) carrying *pfhrp2*-deleted strains hidden in polyclonal infections in the Kenyan samples and one (0.7%) in the Tanzanian samples. Details of the deletions in each country are shown in Table 2.

**Table 2:** Prevalence of *pfhrp2* and *pfhrp3* deletions including in polyclonal infections in clinical samples.

| DNA source | Country | *pfhrp2* and/or *pfhrp3* deletions in samples |  |  | *pfhrp2/3* deletions in polyclonal infections |  |  |
| --- | --- | --- | --- | --- | --- | --- | --- |
|  |  | *pfhrp2* n(%) | *pfhrp3* n(%) | *pfhrp2/3* n(%) | *pfhrp2* n(%) | *pfhrp3* n(%) | *pfhrp2/3* n(%) |
| **Eritrea** | Eritrea (n=50) | 0 | 7 (14) | 32 (64) | 3(0.5) | 4 (8) | 0 |
| **HTD** | Djibouti (n=1) | 0 | 0 | 0 | 0 | 0 | 0 |
|  | Kenya (n=20) | 0 | 0 | 0 | 1 (5) | 1 (5) | 0 |
|  | Eritrea (n=1) | 0 | 0 | 0 | 0 | 0 | 1 (100) |
|  | Ethiopia (n=2) | 0 | 1 (50) | 0 | 0 | 0 | 0 |
|  | Somalia (n=3) | 0 | 0 | 0 | 0 | (2)66.7 | 0 |
|  | South Sudan (n=10) | 0 | 3(30) | 0 | 1 (10) | 0 | 0 |
|  | Sudan (n=29) | 1 (3.5) | 1 (3.5) | 1 (3.5) | 0 | 3 (10.3) | 0 |
|  | Tanzania (n=16) | 0 | 0 | 0 | 0 | 1 (6.3) | 0 |
|  | Uganda (n=27) | 0 | 1 (3.7) | 0 | 2 (7.4) | 3 (11.2) | 0 |
|  | East Africa (n=2) | 0 | 0 | 0 | 0 | 0 | 0 |
| **Kenya/ Tanzania** | Kenya (n=150) | 0 | 0 | 0 | 5 (3.4) | 0 | 0 |
|  | Tanzania (n=149) | 1 (0.7) | 1 (0.7) | 0 | 1 (0.7) | 0 | 0 |

### Estimation of parasite density using *pfldh*

As well as a positive control for parasite DNA quality, the *pfldh* gene was simultaneously used as a parasite target for estimation of parasite density using relative quantification. After assessing its sensitivity, specificity and amplification efficiency the usefulness of the qPCR as an estimator of parasite density was evaluated by comparison to the previously published duplex *pgmet* qPCR assay (18). We performed this comparison using 50 DNA samples from Eritrea and the *pfldh* qPCR showed high degree of correlation (R^2^=0.95) with the duplex *pgmet* qPCR (Figure 3). The *pgmet* qPCR generated relatively higher parasite density compared to the *pfldh* qPCR and this is expected as the former targets multi-copy genes in the apicoplast genome, whereas the latter targets is a single-copy gene.

**Figure 3:**
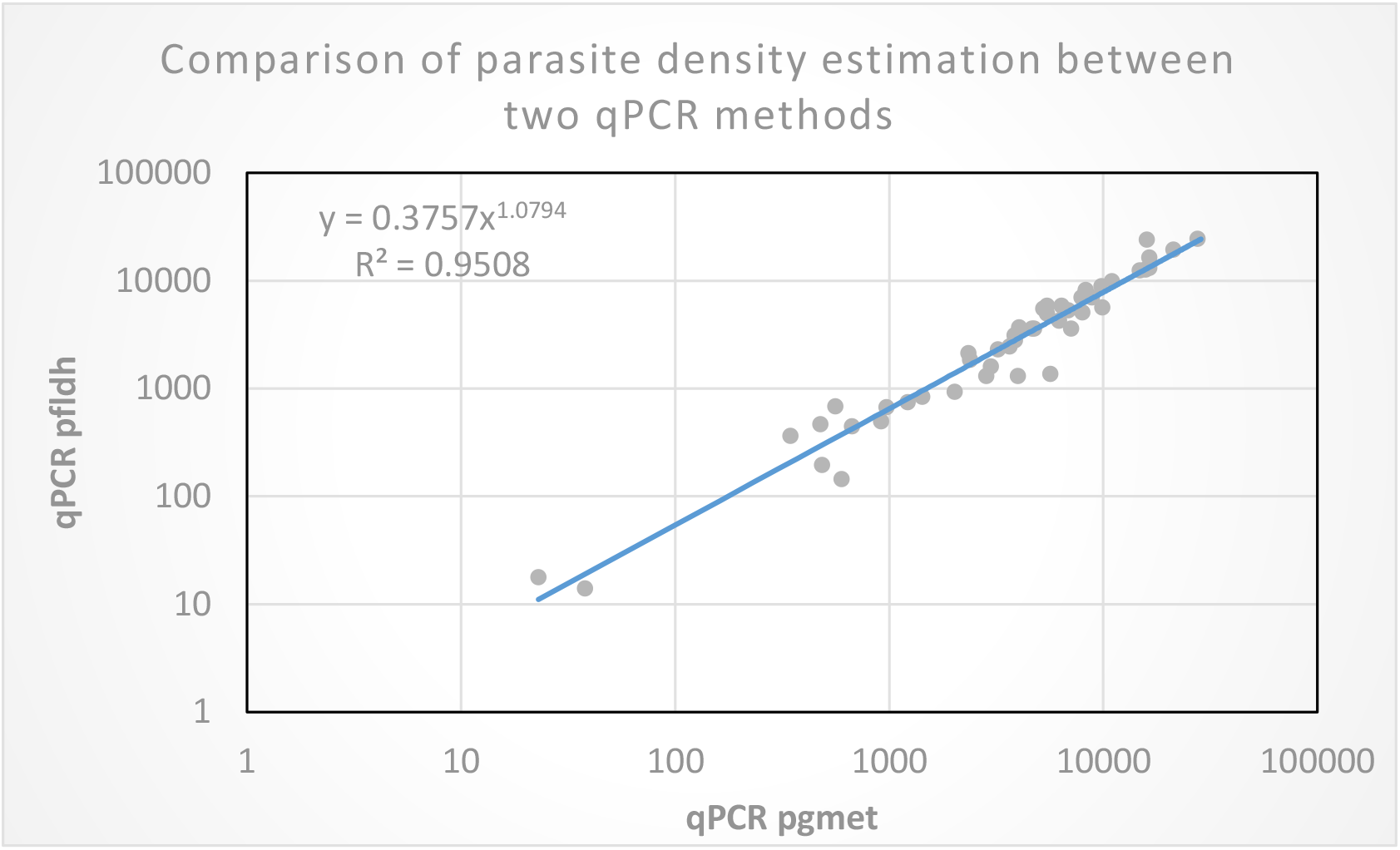
Comparing parasite density estimates produced by *pfldh* and *pgmet* qPCR methods using 50 Eritrean samples. The estimation was done using delta delta relative quantification methods in the presence of *HumTuBB* gene as a normalizer and Pf INT as a calibrator. The two qPCR methods showed strong agreement (R^2^ = 0.95)

## Discussion

In this study, we describe the development, validation and application of a high-throughput multiplex qPCR assay for simultaneous determination of *P. falciparum* with *pfhrp2* and *pfhrp3* deletion genotypes in mono- and polyclonal infections and estimation of parasite density. This highly sensitive and specific method enables accurate and robust assessment of parasite density between 1.5 and 0.76 parasites per μl with a CV of 35% (21). The qPCR detection of each of the targets described here was achieved with high efficiency, R^2^ values greater than 0.96 and very low standard deviation among replicates of each dilution. An additional strength of the assay is the detection of *pfhrp2/3* deletions at relative abundance as low as 20% and as high as 99% in mixed infections, simulated by mixture of cultured parasites. This is particularly relevant in moderate to high transmission settings, where such parasites have already emerged but can be masked in polyclonal infections. The assay can be used as a tool to monitor changes in frequency of deletions providing potential early warning signs of emergence of *P. falciparum* with *pfhrp2/3* deletions. This would allow appropriate measures to be taken to identify, respond and contain the spread of such parasites before they become sufficiently abundant to impact on case management and malaria control programs as has already occurred independently in South America and Eritrea (12, 23).

The design of the multiplex qPCR assay provides several advantages over existing detection methods (5, 14, 24). Firstly, the qPCR assay uses only one parasite target to confirm the presence of DNA and to assess its quality while the gel-electrophoresis based nested PCR (nPCR) methods use three target genes (5, 25, 26). This reduces the cost and throughput time and simplifies the algorithm for interpreting results. Secondly, the parasite target gene (*pfldh*) used for DNA confirmation in the qPCR assay has a single copy, generating equivalent sensitivity to the *pfhrp2/3* targets. When multi-copy genes (e.g., *18SrDNA* and *cytb*) are used in the nPCR and other qPCR assays there is increased risk of false-*pfhrp2/3* deletion calls due to sensitivity differences with the single copy *pfhrp2/3* genes (14, 27). It is difficult to evaluate the accuracy of *pfhrp2/3* deletion calls in the literature, as most laboratories do not report the limit of detection of the nPCR methods used. Thirdly, the qPCR assay uses a human house-keeping gene as internal control. Variability in DNA yield, or loss, introduced during sample collection or parasite DNA extraction or qPCR amplification can thus be corrected for, reducing the risk of false-*pfhrp2/3* deletion calls. Using conventional methods, those samples negative by 18SrDNA may be excluded from further *pfhrp2/3* deletion analysis as they are presumed parasite negatives, potentially underestimating the prevalence of *pfhrp2/3* deletions by missing those samples where technical failure has caused this outcome. Fourthly, the multiplex qPCR assay accurately detects laboratory *P. falciparum* strains and clinical samples with *pfhrp2/3* deletions when mixed as minor or major clones. The ability of the multiplex qPCR assay to detect *pfhrp2/3* deletions in samples with low parasitaemia and in polyclonal infections is made possible due to three unique features: the choice of a single copy parasite gene (*pfldh*) for DNA quality confirmation; the inclusion of a human gene for normalisation and the modification of the primers. Finally, the qPCR assay can also estimate relative parasite density using the human gene as a normalizer and Pf INT as a calibrator. This characteristic of the assay is useful because one of the criteria for confirming deletion of *pfhrp2/3* is estimation of parasite density using microscopy to rule out low parasite density as a factor for lack of parasite target amplification (14). However, microscopy is not always performed during community surveys, and quantification of parasite density using qPCR is required (5, 13). The qPCR assay not only detects *P. falciparum parasites* with *pfhrp2/3* deletions but also simultaneously estimates parasite density in the same experiments, hence reducing time and cost.

Our study shows the presence of *pfhrp2/3* deletions in infected UK travellers for the first time. While *pfhrp2/3* deletions were previously reported in Eritrea (10), Ethiopia (28) and Uganda (29) this is the first time such deletions are reported in Sudan and South Sudan, though a negative *pfhrp2* result was reported in Sudan (30). Interestingly, *pfhrp2* and *pfhrp3* deletions were also detected in polyclonal infections in Kenya, Eritrea, Somalia, South Sudan, Sudan and Uganda. This suggests the circulation of low frequency *pfhrp2*-deleted parasites in Somalia as minor strains in mixed infections. WHO recommends surveillance to determine the prevalence of *pfhrp2/3* deletions occur and in neighbouring areas if the prevalence of *pfhrp2* gene deletions that cause false-negative HRP2-based RDT results in a representative sample is higher than 5%, HRP2-based RDTs should be replaced with alternative *P. falciparum* diagnostic tool that is not exclusively reliant on detection of HRP2 (11). If the prevalence is below 5% a repeat of the survey is recommended in 1-2 years and the detection of *pfhrp2/3* deletions in polyclonal infections by the qPCR assay could be used to inform decisions about how soon to repeat the survey. For example, if the *pfhrp2/3* deletions in polyclonal infections occur in medium to high transmission endemic settings, it may be preferable to survey during the dry season when the multiplicity of infection is lower and would allow accurate estimation of the *pfhrp2/3* deletions.

## Limitations

The challenges of confirming the absence of a gene target require careful attention to lab workflow and DNA quality. First, the sensitivity of the qPCR assay demands careful laboratory workflows that prevent contamination. This is true of all qPCR assays but particularly important for discrimination of low-concentration deleted strains in polyclonal infections. Second, while the qPCR assay was carefully designed and optimized to avoid cross-binding, use of appropriate DNA controls is needed to monitor for unintentional amplification of *pfhrp3* by *pfhrp2* primers and vice versa. We recommend including at least two parasite negative controls (preferably Dd2 and HB3) for *pfhrp2* and *pfhrp3,* respectively, in each experiment. The use of only a double-deleted strain (such as 3BD5) is not recommended. In addition, the qPCR assay targets only one additional single-copy parasite gene while the conventional methods for *pfhrp2* and *pfhrp3* genotyping have employed three independent parasite genes to ensure DNA quality and rule out DNA degradation (14). Because the *pfhrp2* and *pfhrp3* qPCR amplicon lengths are shorter (98bp and 84bp respectively) than typical amplicons generated by the conventional method (~ 300 - 800bp), detection of these targets is expected to be more reliable. If the frequency of deletions is observed to be markedly increased near the limits of detection of the assay in a particular study, then confirmatory testing using a second gene target could be considered. Finally, due to limited conserved regions, the *pfhrp2*-specific primer covered known variant positions present in minority of field samples in the MalariaGEN genome data. The qPCR assay was optimized taking into account the known sequence variants, and the use of nucleotide redundancies in the primer synthesis did not affect the yield in florescence signal (C_q_ value). The variants within our primer sequence are relatively fewer compared to the primer and probe sequences of other recently published qPCR assays (27, 31), which we predict may be challenged by sequence variation (mutations, insertions and deletion) in the primer and probe sequences of *pfhrp2* and *pfhrp3* found in field samples from seven and nine countries respectively (Table S6).

## Conclusion

Our qPCR method for detection of *pfhrp2/3* deletions is a robust alternative with several advantages over existing approaches. The qPCR assay has superior performance to existing methods in speed, cost and ease of interpretation in detecting *pfhrp2/3*-deleted *P. falciparum* parasites from DNA derived from whole blood or filter-paper bloodspots. Data from screening endemic country samples in returning travellers to the United Kingdom suggest systematic surveillance of *pfhrp2/3* deletions in Ethiopia, Sudan and South Sudan is warranted. Careful monitoring of *pfhrp2/3* deletions in Somalia will also be required as the emergence of *pfhrp2/pfhrp3*-deleted parasites as a single-clone infection may soon occur. Based on the findings in this report and elsewhere in the literature, *pfhrp2/3* deletions are present in 31 countries but the scale and scope is still not well elucidated and efforts to dramatically scale up surveillance are needed (32, 33). This qPCR assay can accurately and efficiently support surveillance efforts so that endemic countries have the data required to guide policy on RDT procurement and avert a serious public health threat.

## Declarations

### Author contributions

LG, NS and KBB conceived/designed the study. DN, AB, SM, KM, JJJ, JBP, PC, JC and CJS conceived/designed the individual studies. DAS performed the parasite culturing. JP and SC carried out the genome analysis. LG and RK conducted the initial experiments of the study. KBB conducted the final primer and probe design and analysis, wrote the study protocol and established the assay. LG and KBB conducted the experiments, analysed the results and wrote the first draft. LG, DN, NS, DAS, JBP, PC, JC, CJS, CD and KBB contributed to the reviewing of the manuscript. LG and KB wrote the final manuscript. All authors read and approved the manuscript.

### Ethical approval and consent to participate

All data included in this analysis were obtained in accordance with ethical approvals from country of origin. The data are fully anonymised and cannot be traced back to identifiable individuals. A separate ethical approval was given by London School of Hygiene & Tropical Medicine ethical review committee to conduct this study on the Eritrean samples.

## Acknowledgement

We thank all the patients and staff who participated in this study. We thank Helena Stone for her support with the Malaria Reference Laboratory samples. We also thank Oksana Kharabora for assistance in the lab with the Kenyan and Tanzanian samples. We thank Qin Cheng (ADFMIDI/QIMR Berghofer) sharing with us the *pfhrp2/3* data generated by conventional PCR to validate our qPCR assay.

## Availability of data and materials

Data are available from the corresponding author with reasonable request.

## Consent for publication

Not applicable

## Contributor information

Khalid B Beshir

