## Supplemental Tables and Figures for "A Novel Multiplex qPCR Assay for Detection of *Plasmodium falciparum* with *Histidine-rich Protein 2 and 3 (pfhrp2 and pfhrp3)* Deletions in Polyclonal Infections"

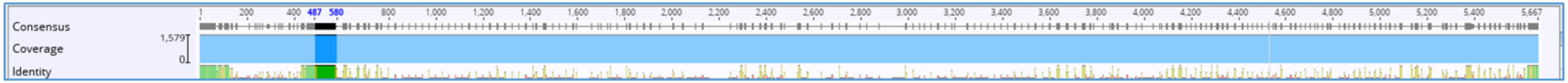


**Figure S1: Location of *pfhrp2* primers and probe: consensus sequence of *pfhrp2* achieved by multiple sequence alignment of 1581 samples obtained from MalariaGEN.**

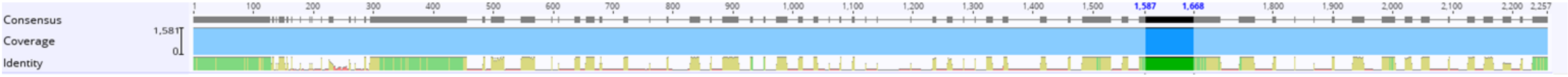


**Figure S2: Location of *pfhrp3* primers and probe: consensus sequence of *pfhrp3* achieved by multiple sequence alignment of 1581 samples obtained from MalariaGEN.**

**Figure 3S: Multiple alignment of *pfhrp2* and *pfhrp3*: The two genes were aligned to check primer cross binding. Two nucleotide (bold) of the 3' end of the *pfhrp2* reverse primer were modified to increase specificity. *Pfhrp2* reverse primer already has one nucleotide difference with *pfhrp3* at the 3' end and this was exploited in the design.**

```
pfhrp2      ATGGTTTCCTTCTCAAAAAATAAAGTATTATCCGCTGCCGTTTTTGCCTCCGTACTTTTG  60
pfhrp3      ATGGTTTCCTTCTCAAAAAATAAAATATTATCCGCTGCCGTTTTTGCTTCCGTACTTTTG  60
            ************************ ********************** ************

pfhrp2      TTAGATAACGTAAGCATTTTAATTGCAAATAGAAATAAATATATCATTCAATTATAAAAA  120
pfhrp3      TTAGATAACGTAAGTATTTTAATTGCAAATATAAATAAATATAACACTTACTTATAAAAA  120
            ************** **************** *********** ** * * *********

pfhrp2      TACATATAATAATAATATATATATATATATATATATATTTATATATTG-----TATATATAT  175
pfhrp3      TACATATAATAATATTATATATATATATATATATATATATATATATATATATATATATATGT  180
            ************** ********************* *******       ******* *

pfhrp2      AAATTTTTTCATTTTTAAATGCTTTTTTATTTTTATATAGAATAATTCCGCATTTAATAA  235
pfhrp3      ACATTTTTACATTTTTAAATG--ATTTTTCATTTTTATAGAATAACTCCGAATTTAACAA  238
            * ****** ***********    ****    *** ********** **** ****** **

pfhrp2      TAACTTGTGTAGCAAAAATGCAAAAGGACTTAATTTAAATAAGAGATTATTACACGAAAC  295
pfhrp3      TAACTTGTTTAGCAAAAATGCAAAAGGACTTAATTCAAATAAGAGATTATTACACGAAAG  298
            ******** ************************** **********************

pfhrp2      TCAAGCACATGTAGATGATGCCCATCATGCTCATCATGTAGCCGATGCCCATCATGCTCA  355
pfhrp3      TCAAGCACATGCAGGTGATGCCCATCATGCACATCATGTAGCTGATGCCCATCATGCACA  358
            *********** ** *************** *********** ************** **

pfhrp2      TCATGTAGCCGATGCCCATCATGCTCATCATGTAGCCGATGCCCATCATGCTCATCATGC  415
pfhrp3      TCATGTAGCTGATGCCCATCATGCACATCATGTAGCTGATGCCCATCATGCACATCATGT  418
            ********* ************** *********** ************** *******

pfhrp2      AGCCGATGCCCATCATGCTCATCATGCAGCCGATGCCCATCATGCTCATCATGCAGCCGA  475
pfhrp3      AGCTGATGCCCATCATGCACATCATGCTCACCATGCAGCTAATGCTCACCATGCAGCTAA  478
            *** ************** ********   * ****   * ******* ********  *

pfhrp2      TGCCCATCATGCTCATCATGCTCACCATGCAGCCGATGCCCATCACGCTCATCATGCAGC  535
pfhrp3      TGCTCACCATGCAGCTAATGCTCACCATGCAGCTAATGCTCATCATGCAGCTAATGCTCA  538
            *** ** *****   * ****************  **** ***** **   * ****

pfhrp2      CGATGCCCATCATGCTCACCATGCAGCTGATGCTCATCACGCTCATCATGCAGCCGATGC  595
pfhrp3      CCATGCAGCTAATGCTCACCATGCAGCTAATGCTCATCATGCAGCTAATGCTCACCATGC  598
            * ****   * ***************** ********** **   * ****   * ****

pfhrp2      CCATCATGCTCACCATGCAGCTGATGCTCATCACGCTCATCATGCATCCGATGCCCATCA  655
pfhrp3      AGCTAATGCTCACCATGCAGCTAATGCTCACCATGCAGCTAATGCTCACCATGCAGCTGA  658
               * ***************** ******* ** **   * ****   * ****   * *

pfhrp2      TGCTCATCATGCAGCCTATGCCCATCATGCTCATCATGCATCCGATGCTCATCATGCAGC  715
pfhrp3      TGCTAATCACGGATTTCATTTTAACCTTCACGATAACAATTCCCATACTTTACATC----  714
            **** **** * *    **    * * *    ** *    *** ** **   ***

pfhrp2      TGATGCTCACCATGCAGCTTATGCCCATCACGCTCATCATGCAGCTGATGCTCATCATGC  775
pfhrp3      --ATGCAAAAGCTAATGCTTGTTTTGATGATTCTCACCATGACGATGCCCACCATGATGG  772
              ****  *   **** *    ** *  **** ****  * **     *** ***

pfhrp2      AGCCGATGCTCACCATGCAACCGATGCTCATCATGCAGCCGATGCTCATCATGCAGCCGA  835
pfhrp3      AGCACACCACGACGATGCCCACCATGA------TGGAG---------CACACCACGACGA  817
            ***  *      ** ****   * ***        ** **            **    * ***

pfhrp2      TGCTCACCATGCAGCCGATGCTCACCATGCAACCGATGCTCATCACGCTCACCATGCAGC  895
pfhrp3      TGCCCACCATGATGGAGCACACCACGACGATGCCCACCATGATGGAGCACACCAC---GA  874
            *** *******  *  *     *** * *   ** *   * **   ** *****    *

pfhrp2      CGATGCTCACCATGCAACCGATGCTCATCACGCTCACCATGCAGCCGATGCTCATCATGC  955
pfhrp3      --------------------------------------------CGATGCCCACCATGA  889
                                                    ****** ** ****

pfhrp2      AGCCGCACACCATGCAACTGATGCTCACCATGCAGCCGCACACCATGCAACCGATGCTCA  1015
pfhrp3      TGGAGCACACCACGATGGAGCAC---ACCACGATGGAGC------------------ACA  928
             *  ******** *     *       **** *  *  **                  **

pfhrp2      CCATGCAGCCGCACACCACGAAGCCGCCACACATTGCCTACGCCATTAAATTTATT      1071
pfhrp3      CCATGATGGAGCACACCATAATGCCACTACGCATCACTTACACCATTAA-------      977
            *****  *  ********  * *** * ** ***  * *** *******
```

*Nucleotide modifications*

*pfhrp2* forward

*pfhrp2* probe

*pfhrp2* reverse

*pfhrp3* forward

*pfhrp3* probe

*pfhrp3* reverse

**Table S1: Primers used for the three parasite target genes and one human gene. *Pfhrp2_R2* primer was modified at 3' end (T->G, highlighted) to increase specificity.. Final primer used in the optimized experiments are highlighted in bold.**

| name | primer sequence | reference |
|---|---|---|
| **Pfhrp2_F1** | 5' TAATTSCGYATTTAATAATAACTTGTG-'3 | This study |
| Pfhrp2_F2 | 5'-TAATTCCGCATTTAATAATAAC**G**TGTG-3' | |
| Pfhrp2_F3 | 5'-TAATTCCGCATTTAATAATAAC**G**TT**G**G-3' | |
| Pfhrp2_R1 | 5'- CATCATCTACATGTGCTTGAG -'3 | |
| **Pfhrp2_R2** | 5'- CATCATCTACATGTGCT**G**GAG -'3 | |
| Pfhrp2_R3 | 5'- CATCATCTACATGTGC**G**TGAG -'3 | |
| **Pfhrp2_probe** | FAM 5'-ATGCAAAAGGACTTAATTTAAATAAGAGATT-'3 BHQ2 | |
| Pfhrp3_F1 | TCCGAATTTAACAATAACTTGTTTAGC | |
| Pfhrp3-R1 | GTCAAGCACATGCAGGTGATG | |
| Pfhrp3_P1 | ATGCAAAAGGACTTAATTCAAATAAGAGATTA | |
| **Pfhrp3_F2** | 5'-TCCGAATTTAACAATAACTTGTTTAGC-'3 | |
| **Pfhrp3_R2** | 5'-GTCAAGCACATGCAGGTGATG-'3 | |
| **Pfhrp3_probe** | JOE 5'-ATGCAAAAGGACTTAATTCAAATAAGAGATTA-'3 BHQ1 | |
| **Pfldh_F** | 5'-ACGATTTGGCTGGAGCAGAT-'3 | Parr *et al*, 2017 |
| **Pfldh_R** | 5'-TCTCTATTCCATTCTTTGTCACTCTTTC-'3 | |
| **Pfldh_probe** | ROX 5'-GTAATAGTAACAGCTGGATTTACCAAGGCCCCA-'3 BHQ1 | |
| **HumTuBB_F** | 5'-AAGGAGGTCGATGAGCAGAT-'3 | Beshir *et al* , 2010 |
| **HumTuBB_R** | 5'-GCTGTCTTGACATTGTTGGG-'3 | |
| **HumanTuBB_P** | CY5 5'-TTAACGTGCAGAACAAGAACAGCAGCT-'3 BHQ2 | |

**Figure S4. Initial experiment using initial *pfhrp2* (left) and *pfhrp3* (right) primers (first panel), after changing *pfhrp3* target sequence (middle panel) and after modification of *pfhrp2* reverse primer (bottom panel)**

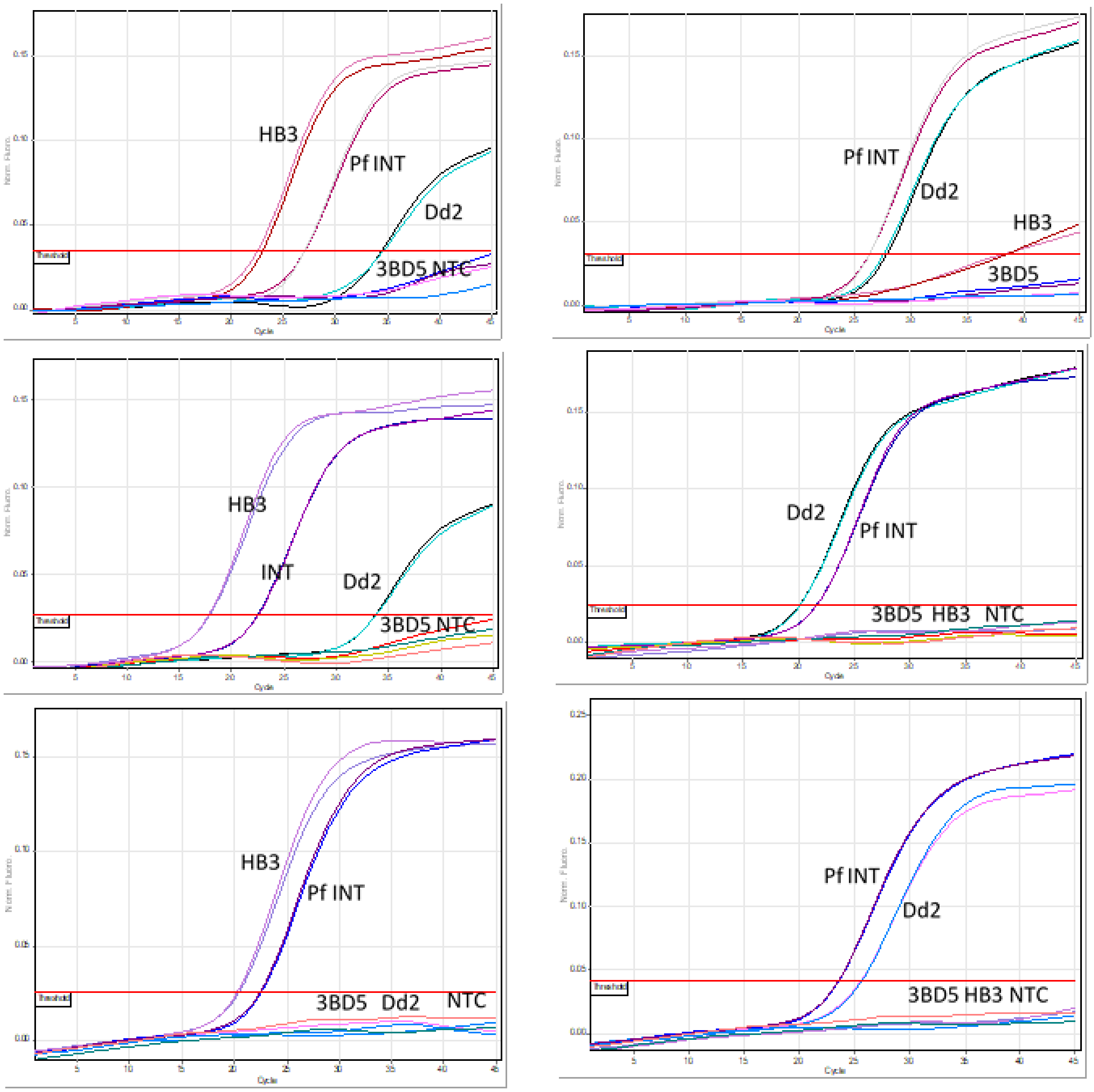

**Table S2. Quantification cycle ($C_q$) produced by different primer combinations. The best $C_q$ value and negative results in 3BD5 and Dd2 was obtained when *pfhrp2*_F1 and *pfhrp2*_R2 were combined. SD, standard deviation.**

| | | Mean $C_q$ value (SD) | | | | | |
|---|---|---|---|---|---|---|---|
| | | **Pfhrp2_F1** | | Pfhrp2_F2 | | Pfhrp2_F3 | |
| | | **$C_q$** | **SD** | **$C_q$** | **SD** | **$C_q$** | **SD** |
| Pfhrp2_R1 | 3BD5 | NEG | | | | NEG | |
| | Dd2 | 42.8 | 0.31 | 32.81 | 0.19 | NEG | |
| | HB3 | 29.5 | 0.11 | 22.54 | 0.13 | 29.15 | 0.71 |
| | pfINT | 30.64 | 0.13 | 23.40 | | 28.04 | 0.17 |
| | NTC | NEG | | NEG | | NEG | |
| **Pfhrp2_R2** | 3BD5 | **NEG** | | NEG | | NEG | |
| | Dd2 | **NEG** | | 34.24 | 0.23 | NEG | |
| | HB3 | **21.08** | 0.06 | 23.53 | 0.50 | 31.35 | 0.64 |
| | pfINT | **22.14** | 0.10 | 24.65 | 0.21 | 30.03 | 0.023 |
| | NTC | **NEG** | | NEG | | NEG | |
| Pfhrp2_R3 | 3BD5 | NEG | | NEG | | NEG | |
| | Dd2 | 33.85 | 0.34 | 31.24 | 0.12 | NEG | |
| | HB3 | 22.17 | 0.12 | 20.98 | 0.38 | 27.25 | 0.04 |
| | pfINT | 23.16 | 0.17 | 22.32 | 0.22 | 27.80 | 0.12 |
| | NTC | NEG | | NEG | | NEG | |

**Table S3. Limit of detection and Limit of quantification of the qPCR assay: the lowest parasite density (3 parasite per µl) positive in the standard dilution was further diluted 2-fold until 0.38 parasite per µl. The lowest diluted sample that can be detected with CV of parasite density of 35% lies between 1.5 and 0.76 parasites per µl.**

| | ***pfhrp2*** | | ***pfhrp3*** | | ***pfldh*** | | ***humTuBB*** | |
|---|---|---|---|---|---|---|---|---|
| **Sample** | **Mean** | **SD** | **Mean** | **SD** | **Mean** | **SD** | **Mean** | **SD** |
| **INT 3 p/ul** | 34.12 | 0.54 | 35.09 | 0.76 | 34.39 | 0.36 | 20.09 | 0.18 |
| **Pf INT 1.5 p/ul** | 36.57 | 0.82 | 36.96 | 0.73 | 36.83 | 0.75 | 19.73 | 0.15 |
| **pf-negative DNA blood** | NEG | | NEG | | NEG | | 20.1 | 0.24 |

**Table S4. Robustness and precision of the qPCR assay: the each assay in the qPCR detected the 8 replicates each of the Pf INT 3 and 1.5 parasites per µl and showed no amplification in the 20 replicates of pf-negative DNA blood.**

| Mean $C_q$ values | | | | | | | |
|---|---|---|---|---|---|---|---|
| Parasite Per µl | ***pfhrp2*** | | ***pfhrp3*** | | ***pfldh*** | | |
| | **Mean** | **SD** | **Mean** | **SD** | **CV** | **Mean** | **SD** |
| 3 | 35.56 | 0.57 | 34.81 | 0.59 | 1.91 | 35.12 | 0.58 |
| 1.5 | 36.77 | 0.80 | 37.28 | 0.71 | 1.90 | 36.69 | 0.74 |
| 0.76 | 38.76 | 1.49 | 38.55 | 1.37 | 3.56 | 38.35 | 1.48 |
| 0.38 | 40.73 | 1.68 | 40.36 | 1.67 | 4.13 | 40.01 | 1.60 |

| Estimated parasite density (parasite per µl) | | | | | | | | | |
|---|---|---|---|---|---|---|---|---|---|
| Parasite Per µl | ***pfhrp2*** | | | ***pfhrp3*** | | | ***pfldh*** | | |
| | **Mean** | **SD** | **CV** | **Mean** | **SD** | **CV** | **Mean** | **SD** | **CV** |
| 3 | 2.86 | 0.24 | 8.34 | 2.95 | 0.29 | 9.81 | 2.62 | 0.23 | 8.83 |
| 1.5 | 0.98 | 0.63 | 32.63 | 0.57 | 0.16 | 26.12 | 0.95 | 0.25 | 28.12 |
| 0.76 | 0.21 | 0.46 | 109.15 | 0.20 | 0.22 | 98.80 | 0.26 | 0.26 | 107.91 |
| 0.38 | 0.05 | 0.12 | 114.77 | 0.06 | 0.07 | 114.14 | 0.07 | 0.08 | 112.26 |

**Table S5. Percentage of *pfhrp2*- and *pfhrp3*-deleted clones in artificially mixed laboratory isolate 3BD5 and Pf INT. The highest and lowest parasite densities in the mixture was 1.5x10$^5$ and 1.5 parasite per µl respectively and this represents the 1:100000 and 100000:1 ratios.**

| **3BD5: Pf INT ratio** | *pfhrp2 deletion %* | | | *pfhrp3 deletion %* | | |
|---|---|---|---|---|---|---|
| | Mean | 95% CI | | Mean | 95% CI | |
| **1:1** | 53.18 | 52.41 | 53.96 | 59.23 | 57.65 | 60.80 |
| **5:1** | 81.90 | 80.84 | 82.97 | 81.31 | 77.08 | 85.55 |
| **10:1** | 91.78 | 91.38 | 92.18 | 92.36 | 90.43 | 94.29 |
| **100:1** | 98.67 | 98.59 | 98.75 | 99.20 | 99.17 | 99.23 |
| **1000:1** | 99.91 | 99.91 | 99.92 | 99.91 | 99.84 | 99.99 |
| **10000:1** | 99.98 | 99.98 | 99.99 | 99.97 | 99.97 | 99.98 |
| **100000:1** | 99.98 | 99.96 | 100.01 | 99.99 | 99.99 | 99.99 |
| **1:1** | 53.92 | 50.86 | 56.97 | 49.65 | 48.52 | 50.77 |
| **1:5** | 17.62 | 9.82 | 25.42 | 18.65 | 16.73 | 20.57 |
| **1:10** | 9.97 | 7.41 | 12.53 | 16.14 | 14.06 | 18.2 |
| **1:100** | 1.06 | 1.01 | 1.12 | 1.29 | 0.78 | 1.80 |
| **1:1000** | 1.07 | 1.00 | 1.15 | 8.05 | 7.81 | 8.30 |
| **1:10000** | 1.86 | 1.56 | 2.16 | 0.14 | 0.06 | 0.22 |
| **1:100000** | 3.27 | 3.16 | 3.38 | 1.29 | 1.14 | 1.43 |

**Table S6: *In silico* analysis of recently published *pfhrp2* and *pfhrp3* qPCR primers and probes**: The sequence analysis was carried out by multiple alignment of *pfhrp2* and *pfhrp3* sequences from 1581 published *P. falciparum* genomes (MalariaGEN) from Africa, SE Asia and south America. The recently published qPCR primers and probes (reference 27 and 31) were searched in the database and mutations (highlighted), insertions (underlined) and deletions (strikethrough) within primer/probe are reported. * Country where mutations were found. ** Both reverse primers showed partial binding to the variable region of *pfhrp2* and *pfhrp3*.

| Reference | Oligo name | Sequence | Country * |
|---|---|---|---|
| Kreidenweiss *et al* (ref. 27 ) | *hrp2* reverse** | GCTACATGATGAGCATGA<br>GCTACATG~~ATGAGCATGA~~<br>GCTACATGGTGAGCATGATGAGCATGA<br>GCTGCATGATGTACATGATGAGCATGA<br>GCTACATGATGGGCATCGGCAACATGATGAGCATGA<br>GCTGCATGATGGGCATCGGCTACATGATGAGCATGA | DRC, Ghana, Laos, Malawi, Thailand and Vietnam |
| | *hrp3* forward** | AGGACTTAATTCAAATAAGAGATTA | Ghana, Guinea, Malawi and Thailand |
| Schindler *et al* (ref. 31 ) | *pfhrp2* forward | GTATTATCCGCTGCCGTTTTTGCC | Ghana |
| | *pfhrp2* reverse | TCTACATGTGCTTGAGTTTCG | Bangladesh, Cambodia, Laos and Vietnam |
| | *pfhrp2* probe | TTCCGCATTTAATAATAACTTGTGTAGC | The Gambia, Ghana and Mali |
| | *pfhrp3* forward | ATATTATCCGCTGCCGTTTTTGCT | Malawi |
| | *pfhrp3* probe | CTCCGAATTTAACAATAACTTGTTTAGC | Bangladesh, Ghana, Guinea, Malawi and Mali |
